# Multi-omics biomarker pipeline reveals elevated levels of protein-glutamine gamma-glutamyltransferase 4 in seminal plasma of prostate cancer patients

**DOI:** 10.1101/120873

**Authors:** Andrei P. Drabovich, Punit Saraon, Mikalai Drabovich, Theano D. Karakosta, Apostolos Dimitromanolakis, M. Eric Hyndman, Keith Jarvi, Eleftherios P. Diamandis

**Affiliations:** Department of Laboratory Medicine and Pathobiology, University of Toronto, Toronto, Ontario, M5T 3L9 Canada; Department of Clinical Biochemistry, University Health Network, Toronto, Ontario, M5T 3L9 Canada; Department of Pathology and Laboratory Medicine, Mount Sinai Hospital, Toronto, Ontario, M5T 3L9 Canada; Independent researcher, Palo Alto 94303, USA; Prostate Cancer Centre, Southern Alberta Institute of Urology, Rockyview General Hospital, Calgary, Alberta, T2V 1P9, Canada; Lunenfeld-Tanenbaum Research Institute, Mount Sinai Hospital, Toronto, Ontario, M5T 3L9 Canada; Department of Surgery, Division of Urology, Mount Sinai Hospital, University of Toronto, Toronto, Ontario, M5T 3L9 Canada

**Keywords:** prostate cancer, biomarker discovery, seminal plasma, proteomics, mass spectrometry, selected reaction monitoring, TGM4, protein-glutamine gamma-glutamyltransferase 4

## Abstract

**Purpose:** Prostate-specific antigen, a blood serum biomarker of prostate cancer, lacks specificity and prognostic significance, so considerable efforts are devoted to developing novel biomarkers. Seminal plasma, due to its proximity to prostate, is a promising fluid for biomarker discovery and non-invasive diagnostics. In this study, we investigated if seminal plasma proteins could increase specificity of detecting primary prostate cancer and discriminate between high- and low-grade cancers.

**Experimental Design:** To select 148 most promising biomarker candidates, we combined proteins identified through five independent data mining or experimental approaches: tissue transcriptomics, seminal plasma proteomics, cell secretomics, tissue specificity and androgen regulation. A rigorous biomarker development pipeline based on targeted proteomics assays was designed to evaluate the most promising candidates.

**Results:** We qualified 77 and verified 19 proteins in seminal plasma of 67 negative biopsy and 155 prostate cancer patients. Verification revealed a prostate-specific, secreted and androgen-regulated protein-glutamine gamma-glutamyltransferase 4 (TGM4), which could predict prostate cancer on biopsy and outperformed age and serum PSA. Machine-learning approaches also revealed improved multi-marker combinations for diagnosis and prognosis. In the independent verification set measured by an inhouse ELISA, TGM4 was up-regulated 3.7-fold (*P*=0.006) and revealed AUC 0.66 for detecting prostate cancer on biopsy for patients with serum PSA≥4 ng/mL and age≥50. Low levels of TGM4 (median 120 pg/mL) were detected in blood serum, but could not differentiate between negative biopsy, prostate cancer or prostate inflammation.

**Conclusions:** Performance of TGM4 warrants its further investigation within the distinct genomic subtypes and evaluation for the inclusion into emerging multi-biomarker panels.

## Introduction

Prostate cancer is the most frequently diagnosed neoplasm and the third leading cause of cancer mortality in men. Its incidence rate has continued to increase rapidly during the past two decades, especially in men over the age of 50 years. Worldwide, close to 260,000 men die from prostate cancer every year (1). Our best current strategy to help prostate cancer patients is early diagnosis and administration of the most appropriate therapy, including active surveillance only (2).

The most commonly used prostate cancer biomarker, prostate-specific antigen (PSA), is secreted by both normal prostate cells and prostate cancer cells. There is no question that the introduction of PSA testing over the last two decades revolutionized the practice of urology. As a result of PSA screening, most men today with prostate cancer present with localized disease and serum PSA values <10 ng/mL. However, the widespread use of PSA screening is not without controversy (3,4).

Although PSA is an excellent biochemical marker, it has a number of important limitations, including a lack of specificity and a lack of prognostic significance. PSA expression is prostatic tissue-specific but not prostate cancer-specific. Serum PSA levels are increased in both prostate cancer and in other non-malignant prostatic diseases, including benign prostatic hyperplasia and prostate inflammation. Because of the above limitations, clinicians currently perform on average four prostatic biopsies in order to detect one prostate cancer. PSA levels also do not predict the clinical significance or aggressiveness of prostate cancer. Most men with prostate cancer are destined to die of another condition before prostate cancer becomes clinically significant (5). Lack of specificity and prognostic significance are two major limitations of PSA and constitute the major unmet needs in the current clinical diagnostics of prostate cancer.

There have been intense efforts for the identification of novel prostate cancer biomarkers in blood or urine. Most promising blood serum-based biomarkers included human kallikrein 2 (6) and different forms of PSA (7). A few of the more promising RNA biomarkers in urine included PCA3 (8), TMPRSS2-ERG fusion (9) and SPINK1 (10). Of these, FDA-approved tests include PCA3 in urine to aid in decisions for the second biopsy (11) and Prostate Health Index (7) in blood of men 50 years and older with elevated PSA and negative digital rectal examination. None of novel biomarkers in blood or urine has as yet proven to be significantly more sensitive or specific than PSA or predicted prostate cancer aggressiveness.

There is little or no reported work on the study of prostate cancer biomarkers in seminal plasma (SP), even though much of the work to identify and characterize PSA was originally carried out in SP (12). Nearly a quarter of molecular composition of SP is secreted by prostate (13), with the rest produced by seminal vesicles, epididymis, testis and periurethral glands. We previously completed several years of work on proteome of SP and identified more than 3,000 proteins in SP of healthy men and patients with infertility, prostatitis and prostate cancer (14,15). Our work resulted in first-of-a-kind SP biomarkers for the differential diagnosis of male infertility (16–18). Success with male infertility biomarkers motivated us to apply a similar strategy to prostate cancer.

In this work, we hypothesized that some proteins expressed by the prostate and secreted into SP can predict PCa on biopsy or differentiate between low- and high-grade PCa. To select the most promising biomarker candidates, we combined proteins identified through five data mining and experimental - omics approaches, such as transcriptomics, proteomics, secretomics, tissue specificity and androgen regulation. Only those proteins which were previously identified in the SP proteome were considered as candidates and were verified by mass spectrometry-based selected reaction monitoring (SRM) assays. Powerful nonlinear machine-learning algorithms (19) were utilized to evaluate multi-marker models for prostate cancer diagnosis and prognosis. Our study was designed to simultaneously assess biomarker candidates for the two clinical needs: (i) differentiation between PCa and negative biopsy, and (ii) discrimination between low- and high-grade PCa. To our knowledge, this work is the largest and the most comprehensive study on seminal proteins in prostate cancer.

## Materials and methods

### Study design

The objectives of this study were to select the most promising PCa biomarker candidates, develop quantitative mass spectrometry assays, qualify the most promising candidates, verify candidate biomarkers in a large set of SP samples by mass spectrometry and verify top protein by ELISA in SP and blood serum samples of patients with primary PCa and benign disease.

### Patients and clinical samples

SP samples with relevant clinical information were obtained through the Murray Koffler Urologic Wellness Centre at Mount Sinai Hospital (REB #08-0117-E), University Health Network (#09-0830-AE) and Calgary Prostate Cancer Center (#18166). Men referred for a prostate biopsy were asked to participate in this study. None of these men had clinical signs of prostate inflammation. Following prostate biopsy, men were categorized as having: (i) low-grade PCa (Gleason Score, GS, =6, median serum PSA 5 ng/mL [IQR 4-8 ng/mL], median age 63 y.o. [IQR 57-67 y.o.], N=125); (ii) intermediate-grade PCa (GS=7, 3+4, median serum PSA 6 ng/mL [IQR 5-7 ng/mL], median age 62 y.o. [IQR 56-67 y.o.], N=50); (iii) high-grade PCa (GS=7, 4+3 and ≥ 8; median serum PSA 9 ng/mL [IQR 7-11 ng/mL], median age 65 y.o. [IQR 60-70 y.o.], N=25); (iv) no evidence of cancer (negative biopsy, median serum PSA 6 ng/mL [IQR 4-7 ng/mL], median age 60 y.o. [IQR 55-64 y.o.], N=68). Semen samples were collected by masturbation into a sterile collection cup either at home or at urology clinics. Following liquefaction for 1 hour at room temperature, semen samples were centrifuged at 13,000 g for 15 min, and the supernatants were frozen at −80°C. Blood serum samples included: (i) confirmed primary PCa (all GS scores, median serum PSA 9 ng/mL [IQR 6-16 ng/mL], median age 65 y.o. [IQR 55-74 y.o.], N=29); (ii) no evidence of cancer (negative biopsy, median serum PSA 8 ng/mL [IQR 6-12 ng/mL], median age 63 y.o. [IQR 54-69 y.o.], N=23); (iii) prostate inflammation (median serum PSA 8 ng/mL [IQR 6-14 ng/mL], median age 65 y.o. [IQR 56-70 y.o.], N=11); (iv) healthy men (median age 36 y.o. [IQR 31-41 y.o.], N=17). Blood samples were collected at the diagnostic laboratory at Mount Sinai Hospital, and blood serum was stored at −80°C.

### Differential proteomics

Mass spectrometry was used to identify differentially expressed proteins in three pools of SP samples from negative biopsy, low-grade PCa and high-grade PCa patients. SP pools included: (i) low-grade PCa (GS=6, median serum PSA 8 ng/mL [IQR 6-10 ng/mL], median age 65 y.o. [IQR 62-67 y.o.], N=5); (ii) high-grade PCa (GS=8 or 9; median serum PSA 14 ng/mL [IQR 9-19 ng/mL], median age 66 y.o. [IQR 66-66 y.o.], N=5); (iii) no evidence of cancer (negative biopsy, median serum PSA 7 ng/mL [IQR 6-10 ng/mL], median age 63 y.o. [IQR 55-65 y.o.], N=5). The rationale for using pooled samples was to reduce the effects of the protein biological variability between patients and to increase the likelihood of identifying protein differences which would be consistent between patients with a similar diagnosis. Each pool was subjected to the proteomic sample preparation and mass spectrometry analysis in triplicates (whole process replicates). Tryptic digestion of proteins (500 μg total protein per pool) was performed as previously described (16,20). Briefly, proteins were denatured at 65°C in the presence of 0.02% of RapiGest SF (Waters, Milford, MA), reduced with 10 mM dithiothreitol, alkylated with 20 mM iodoacetamide and digested overnight at 37°C using sequencing grade modified trypsin (trypsin:total protein ratio 1:30; Promega, Madison WI, USA). RapiGest SF was cleaved with 1% trifluoroacetic acid and removed by centrifugation. Following protein digestion, peptides were fractionated by strong-cation exchange chromatography, twenty three fractions were collected for each pool replicate, concentrated with 10 μL OMIX C18 tips (Varian, Lake Forest, CA) and analyzed by the reverse phase liquid chromatography-tandem mass spectrometry (LTQ-Orbitrap XL, Thermo Scientific), as previously described (21). XCalibur software (v. 2.0.5; Thermo Scientific) was used to generate RAW files. MaxQuant software (version 1.1.1.25) was used for protein identification and label-free quantification (LFQ). MaxQuant executed spectral search against a concatenated International Protein Index (IPI) human protein database (version 3.71) and a decoy database. Parameters included: trypsin enzyme specificity, 1 missed cleavage, minimum peptide length of 7 amino acids, minimum of 1 unique peptide, top 6 MS/MS peaks per 100 Da, peptide mass tolerance of 20 ppm for precursor ion and MS/MS tolerance of 0.5 Da and fixed modification of cysteines by carbamidomethylation. Variable modifications included oxidation of methionine and acetylation of the protein at N-terminus. All entries were filtered using a false positive rate of 1% both at the peptide and protein levels, and false positives were removed. MaxQuant search file proteinGroups.txt was uploaded to Perseus software (version 1.4.1.3). Protein identifications annotated in the columns “Only identified by site”, “Reverse” and “Contaminant” as well as proteins identified only in a single replicate were filtered out. LFQ intensities were log2-transformed and used to calculate means and statistical significance (t-test with Benjamini-Hochberg false-discovery rate-adjusted P-values) and generate volcano plots. Raw mass spectrometry proteomics data were deposited to the ProteomeXchange Consortium via the PRIDE partner repository with the dataset identifier PXD007657. Reviewer account details (http://www.ebi.ac.uk/pride/archive/login): Username: Password: pyciV5y7.

### Development of SRM assays for the qualification phase

SRM assays were developed as previously described (22–24). Briefly, the Peptide Atlas (www.peptideatlas.org) was used to select top 5-7 peptides (charge +2 and +3) for each of 148 candidate proteins and 11 control proteins (representing other six glands or cell types in the male urogenital system). Fully tryptic peptides with 7-20 amino acids were chosen, and peptides with methionine and N-terminal cysteine residues were avoided, if possible. A list of peptides and top 7 transitions were downloaded. All proteins were ranked according to their MaxQuant LFQ intensities and split into groups of high-, medium- and low-abundance SP proteins. Sixty survey methods with 15 peptides, 7 transitions per peptide, 20 ms scan times, 8 minute scheduling windows based on predicted retention times were designed and tested with multiple iterations in the digest of normal SP. The rationale for multiple iterations was to quickly develop SRM methods for high-abundance peptides and exclude them from the subsequent iterations, thus focusing on medium- and low-abundance proteins. Nearly 900 peptides and 6,000 transitions were experimentally tested on TSQ Vantage in the digest of normal SP. Raw files were uploaded to Pinpoint software (Thermo Scientific), and peaks were analyzed manually. High-abundance peptides with clear peaks, high signal-to-noise intensities and multiple overlapping transitions were selected, while medium- and low-abundance peptides moved to the second iteration. Peptides or transitions in doubt were confirmed with our SP proteome data. In the second iteration, we designed 37 methods (~2500 transitions) and tested them with 35 ms scan times, thus lowering background and facilitating detection of low-abundance proteins. In the third iteration, we tested 9 methods (~500 transitions) with 40 ms scan times. In the fourth iteration, we experimentally reconfirmed all peptides and verified, recorded or optimized the following parameters: (i) top 3 transitions; (ii) retention times and scheduling intervals; (iii) selectivity of transitions and possible interferences; and (iv) scan times. Transitions with fragment m/z higher than precursor m/z were preferable; however, some transitions with lower m/z but high signal-to-noise ratio were also used. For proteins with multiple peptides, a single peptide with the highest SRM area was chosen. All peptides were analyzed with the Basic Local Alignment Search Tool at http://blast.ncbi.nlm.nih.gov/Blast.cgi to ensure that peptides were unique to each UniProtKB/Swiss-Prot protein identifier. In the final iteration, 105 candidate and control proteins (107 peptides, 321 transitions) were scheduled in a single SRM method within 2.8-min (±1.4 min) intervals during a 60 min LC gradient. Three most intense and reproducible transitions were monitored per each peptide. Scan times were optimized for each peptide to ensure acquisition of 15-20 points per LC peak per transition and varied between 4 and 25 ms.

### Relative quantification by label-free SRM in the qualification phase

Ten microliters of each SP were diluted 10-fold with 50 mM ammonium bicarbonate (pH 7.8), and an aliquot equivalent to 0.5 μL of SP was subjected to the proteomic sample preparation. SP samples were randomized on a 96-well plate and included: (i) low-grade PCa (GS=6, median serum PSA 6 ng/mL [IQR 5-8 ng/mL], median age 64 y.o. [IQR 62-68 y.o.], N=24); (ii) high-grade PCa (GS=7, 4+3 and ≥ 8; median serum PSA 10 ng/mL [IQR 7-12 ng/mL], median age 67 y.o. [IQR 64-71 y.o.], N=14); (iii) no evidence of cancer (negative biopsy, median serum PSA 6 ng/mL [IQR 5-7 ng/mL], median age 63 y.o. [IQR 59-65 y.o.], N=13). Proteins were denatured at 60°C with 0.1% Rapigest SF, and the disulfide bonds were reduced with 10 mM dithiothreitol. After reduction, the samples were alkylated with 20 mM iodoacetamide. Samples were then trypsin-digested overnight at 37°C. One hundred and eighty femtomoles of heavy isotope-labeled peptide LSEPAELTDAVK of KLK3 were spiked into each digest and used as a quality control for C18 microextraction and data normalization. Rapigest was cleaved with 1% trifluoroacetic acid, and a 96-well plate was centrifuged at 2,000 rpm for 20 min. Each digest was subjected to micro extraction with 10 μL OMIX C18 tips (Varian, Inc.). The plate was stored at −20°C and thawed prior to SRM. Each sample was analyzed by SRM in triplicates. One to four mass spectrometry quality control samples were run every 12 injections. The LC EASY-nLC 1000 (Thermo Fisher Scientific Inc.) was coupled online to TSQ Vantage triple-quadrupole mass spectrometer (Thermo Fisher Scientific Inc.) using a nanoelectrospray ionization source. Peptides were separated on a 2 cm trap column (150 μm ID, 5 μm C18) and eluted onto a 5 cm resolving column (75 μm ID, 3 μm C18). Peptides were scheduled within 2.8-min intervals during a 60 min LC gradient. SRM method had the following parameters: optimized collision energy values, 0.010 m/z scan width, 4 – 25 ms scan times, 0.4 FWHM resolution of the first quadrupole (Q1), 0.7 FWHM resolution of the third quadrupole (Q3), 1.5 mTorr pressure of the second quadrupole, tuned S-lens values, +1 V declustering voltage. Carryover was estimated in the range 0.05-0.2%. Raw files recorded for each sample were analyzed using Pinpoint software. Areas of all peptides were normalized to the spike-in standard heavy isotope-labeled peptide LSEPAELTDAVK of KLK3 and used with GraphPad PRISM (version 5.03) to calculate ROC AUC areas, sensitivities and specificities.

### Upgrade of SRM assays for the verification phase

Purified heavy isotope-labeled peptides (SpikeTides ™_TQL) with trypsin-cleavable quantifying tags (serine-alanine-[3-nitro]tyrosine-glycine) and quantified amounts (1 nmol per aliquot) were obtained from JPT Peptide Technologies GmbH, Berlin, Germany. SRM transitions and TSQ Vantage instrumental parameters were validated, optimized or corrected using high concentrations of digested synthetic peptides. Retention times and scheduling windows were optimized for a 30 min LC gradient. Synthetic peptides were also used to assess the efficiency of digestion of peptide internal standards by trypsin and assess chemical modifications of peptides (cysteine alkylation, methionine oxidation, formation of pyroglutamate of N-terminal glutamine and deamidation of asparagines and glutamines) (25). Upgraded and optimized SRM assay for the verification phase was used to assess numerous pre-analytical variables for each protein using three SP samples: LC-SRM injection reproducibility, trypsin digestion reproducibility, impact of different total protein concentrations in SP samples and day-to-day reproducibility of the whole process of sample processing and analysis.

### Absolute quantification by SRM in the verification phase

SP samples (N=219) were randomized between six 96-well plate and included: (i) low-grade PCa (GS=6, median serum PSA 5 ng/mL [IQR 4-7 ng/mL], median age 64 y.o. [IQR 59-68 y.o.], N=97); (ii) intermediate-grade PCa (GS=7, 3+4; median serum PSA 6 ng/mL [IQR 5-7 ng/mL], median age 62 y.o. [IQR 55-66 y.o.], N=36); (iii) high-grade PCa (GS=7, 4+3 and ≥ 8; median serum PSA 9 ng/mL [IQR 7-12 ng/mL], median age 67 y.o. [IQR 63-71 y.o.], N=19); (iii) no evidence of cancer (negative biopsy, median serum PSA 6 ng/mL [IQR 4-7 ng/mL], median age 60 y.o. [IQR 55-65 y.o.], N=67). SP samples were processes and analyzed in the same way as in the qualification phase, except the following:

(i) Six hundred femtomoles of heavy isotope-labeled peptides (SpikeTides ™_TQL) were spiked into 5 μL of 10-fold diluted SP (~20 μg of total protein) prior to sample preparation and trypsin digestion;
(ii) Peptides were measured by TSQ Quantiva (Thermo Scientific, San Jose, CA), as previously described (26). Polarity was set to “positive”, ion transfer tube temperature was 300°C, CID argon pressure was 2.0 mTorr, and source fragmentation was 10 volts. The resolution settings of the first and third quadrupoles were 0.2 and 0.7 Da FWHM, respectively. High resolution in the first quadrupole facilitated exclusion of potentially interfering ions, thus improving selectivity in the complex biological matrices. Dwell times varied between 4 and 40 ms, depending on protein abundance in SP. Thirty seven peptides, 82 parent ions (including light and heavy forms as well as additional +3 forms for CORO1B, GALNT7, PROS1 and an N-terminal pyroglutamate form for STEAP4) and 250 transitions were scheduled within 2.2-min intervals during a 30 min LC gradient. Each sample was analyzed by SRM in duplicates.
(iii) Each of six plates included its own set of calibration curves which was generated by spiking serial dilutions of heavy internal standard peptides (0.4, 4, 40, 400, 4000 and 12000 pmol/mL) into a single SP sample digest. Non-linear regression point-to-point fit curves with 20,000 points were calculated in GraphPad PRISM (version 5.03) within 0.4-12000 pmol/mL concentration ranges.
(iv) Areas of all peptides were normalized to the corresponding heavy peptide internal standards. Normalized areas were fitted to corresponding calibration curves for each plate, and the absolute concentrations of each protein in each sample were calculated. Average concentration of three injections was calculated. Molecular masses calculated from the full canonical sequences without post translational modifications according to the UniProtKB database were used to translate concentrations from pmol/mL to μg/mL. SRM raw mass spectrometry data for both phases were deposited to the Peptide Atlas repository (http://www.peptideatlas.org/PASS/PASS00989) with the dataset identifier PASS00989 and the following credentials: Username: PASS00989; Password: BU5655ot.

### Machine-learning analysis

Scikit-learn library for the Python programming language was used for machine learning analysis (27). Multiple algorithms including generalized linear models, support vector machines, nearest neighbors, naive Bayes, shallow and deep neural networks were tested. A nonlinear Extreme Gradient Boosting (XGBoost) algorithm (19) was selected as the most efficient approach for selection of weak features with small data sets and for generation of a single strong classifier with a set of weak classifiers. XGBoost was also relatively fast and allowed for even further speed improvements via standard parallelization strategies for multi-processor systems.

Briefly, twenty two variables (SP proteins, serum PSA and age) were used to generate all possible 1-, 2-, 3-, 4- and 5- marker combinations (35,442 combinations in total). Cross-validation was run for each combination in order to select combinations with the highest *F*_05_-scores. In comparison to *F*_1_-score with the harmonic mean of precision (or positive predictive value) and recall (or sensitivity), *F*_05_-score weighted precision twice as much as recall. AUCs, sensitivities, specificities and negative predictive values were also estimated. Over-fitting was reduced using stringent 10×10 cross-validation which included splitting data 10 times into 10 sets in a way to keep the proportion of negative and positive samples across all sets approximately the same as in the whole dataset. The stratified 10-fold cross-validation was repeated 10 times, and the total number of train-validation runs was 100. Top combinations were verified on the whole dataset of patients to ensure that each potential marker had feature scores higher than a randomly generated feature. Finally, 100-fold bootstrapping was used to estimate mean values for performance metrics and calculate 95% confidence intervals.

### Development of TGM4 ELISA

Anti-TGM4 polyclonal mouse antibodies generated against the full length TGM4 (Met1-Lys684) were obtained from Abnova (H00007047-B01P; lot 09177 WUIZ). Antigen affinity-purified anti-TGM4 polyclonal sheep antibodies generated against the full length TGM4 (Met2-Lys684) were obtained from R&D Systems (AF5760; lot CDCX0111121). The full length recombinant human TGM4 Met2-Lys684 protein (R&D Systems; AAC50516; lot SIC0213111) expressed in S. frugiperda insect ovarian cell line Sf 21 was used during assay development. SP pool with a known concentration of endogenous TGM4 (4.65 μg/mL), as measured by SRM, was used as the assay calibrator.

To validate specificity of antibodies, we first performed immunocapture-SRM assays with recombinant human (rhTGM4) and endogenous TGM4 from SP. ELISA plates were coated with 500 ng/well of either sheep or mouse antibodies and incubated with either rhTGM4 (0, 2, 10 and 51 ng/mL) or endogenous TGM4 (0, 4, 19 and 94 ng/mL). Following that, plates were washed, proteins were digested with trypsin, and peptides were quantified with an SRM assay. As a result, endogenous TGM4 from SP was enriched equally well by both antibodies, while sheep antibody AF5760 could not efficiently capture rhTGM4.

We then coated ELISA plates with 300 ng/well of either sheep or mouse antibodies, incubated with either rhTGM4 (0, 5, 20 and 100 ng/mL) or endogenous TGM4 in SP (0, 5, 20 and 100 ng/mL) and detected with biotinylated sheep or mouse antibodies and standard time-resolved fluorescence ELISA format (18). ELISA revealed that endogenous TGM4 from SP generated much higher signal than rhTGM4, and that the mouse-sheep format generated substantially lower background and higher signal-to-noise. Finally, mouse-sheep format with endogenous TGM4 as a calibrator was selected due to higher signal-to-noise ratio. Limit of blank and limit of detection of our in-house ELISA were 9 and 22 pg/mL, respectively.

To measure TGM4 in SP and blood serum samples, the 96-well ELISA plates were coated with 300 ng/well of mouse antibody H00007047-B01P in 50 mM Tris-HCl buffer at pH 7.8. Plates were washed twice with the washing buffer (0.05% Tween 20 in 20 mM Tris-HCl and 150 mM NaCl at pH 7.4). Calibrator and SP samples were diluted with the assay diluent (60 g/L BSA, 25 mL/L normal mouse serum, 100 mL/L normal goat serum and 10 g/L bovine IgG in 50 mM Tris-HCl at pH 7.8). Serial dilutions of the calibrator (SP pool with 4.65 μg/mL endogenous TGM4 diluted to 0.02, 0.06, 0.18, 0.56, 1.7, 5 and 15 ng/mL, 100 μL/well) were prepared. Patient SP samples (2 μL) were diluted 400-fold with the assay diluent and added on ELISA plates (100 μL/well). Following 2 hour incubation with gentle shaking, plates were washed twice with the washing buffer. Sheep polyclonal antibody AF5760 was biotinylated in-house, added to each well (25 ng in 100 μL of the assay diluent) and incubated for 1 hour. Plates were washed six times, and streptavidin-conjugated alkaline phosphatase was added for 15 min with gentle shaking. After the final wash, diflunisal phosphate solution was prepared in the substrate buffer (0.1 M NaCl, 1 mM MgCl_2_ in 0.1 M Tris at pH 9.1), added to the plate (100 μL per well) and incubated for 10 min at room temperature with gentle shaking. Finally, the developing solution (1 M Tris-HCl, 0.4 M NaOH, 2 mM TbCl_3_ and 3 mM EDTA) was added and mixed for 1 min. Time-resolved fluorescence was measured with the Wallac EnVision 2103 Multilabel Reader (Perkin Elmer), as previously described(18). All SP and blood serum samples were measured in duplicates. SP samples with TGM4 concentrations outside the range were re-measured with 800-, 40- and 10-fold dilutions. Blood serum samples (50 μL each) were analyzed with 2-fold dilutions.

## Results

### Selection of candidate proteins

We combined transcripts or proteins identified through five independent data mining or experimental approaches and generated a list of the most promising 148 biomarker candidates (**Fig. 1A**). To facilitate our diagnostic strategy, we considered as candidates only secreted and membrane-bound proteins which were previously identified in our SP proteome of more than 3,000 proteins (14,15,28). We assumed that some candidates selected with large-scale -omics approaches might be false-positives due to various pre-analytical, analytical and data analysis biases. We also assumed that different -omics approaches may have certain limitations (for example, discrepancy between mRNA and protein fold changes or inability to measure low-abundance SP proteins by mass spectrometry). As a result, our candidates were not compared across all five approaches, but were independently selected and merged into a single list. Our strategy was to apply more relaxed criteria for the selection of candidates, but then perform very stringent verification of candidates in SP by high-quality quantitative assays (**Fig. 1B**).

**Fig. 1.**
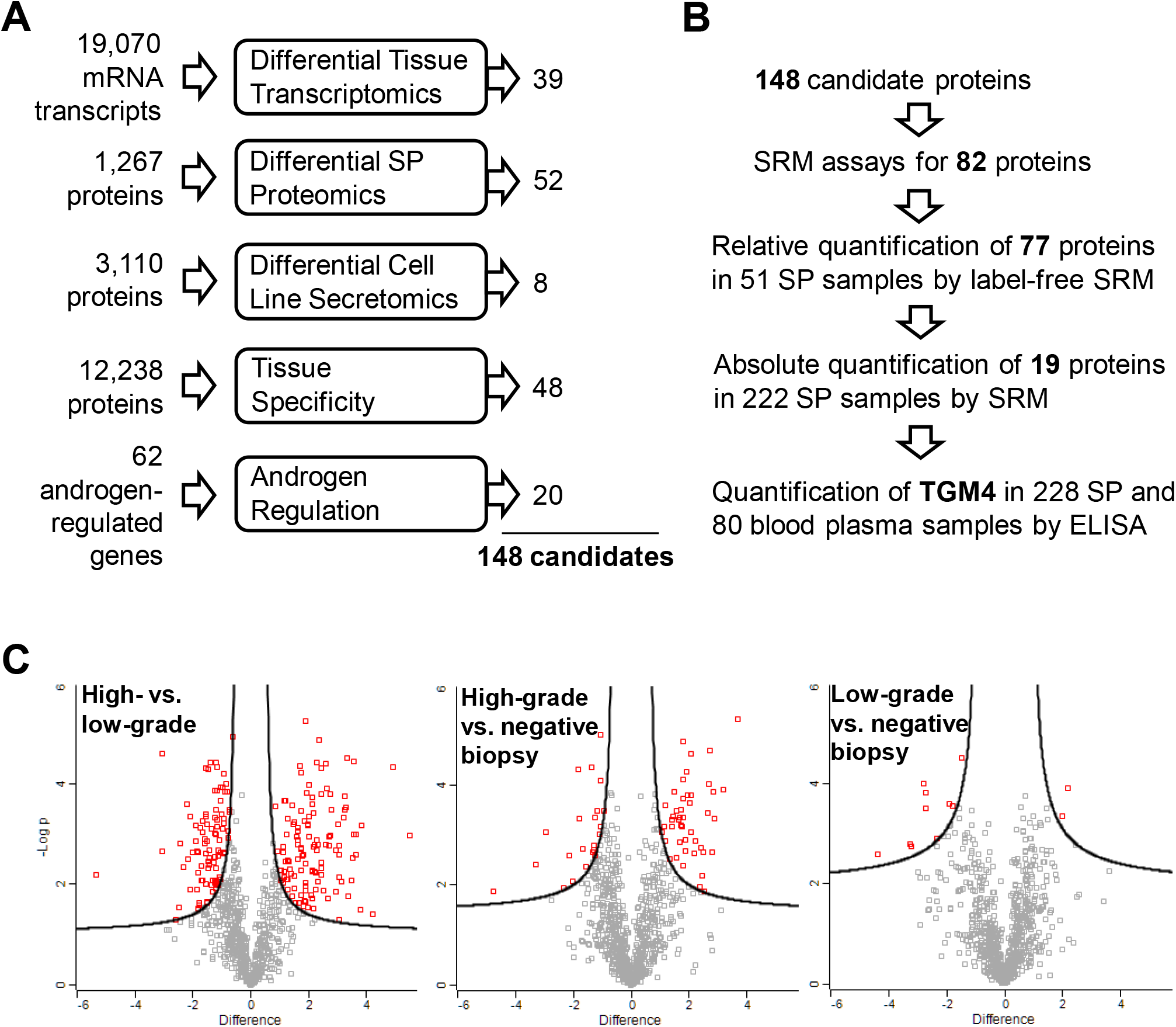
Selection of candidate proteins. (**A**) The most promising PCa biomarker candidates were generated through five independent experimental and data mining approaches. (**B**) The combined 148 candidates were subjected to the SRM method development, followed by qualification and verification in SP and blood serum. (**C**) A bottom-up proteomic approach and two-dimensional liquid chromatography followed by shotgun mass spectrometry and label-free quantification were used to identify differentially expressed proteins in pools of SP samples from patients with negative biopsy (NBx, serum PSA >4 ng/mL, N=5), low-grade PCa (LG, GS =6, PSA >4 ng/mL, N=5) and high-grade PCa (GS ≥8, PSA >4 ng/mL, N=5) patients. Log2 differences and the t-test with Benjamini-Hochberg false-discovery rate-adjusted P-values were calculated with Perseus software, and 1% FDR was used as a cut-off to select differentially expressed protein for each comparison. High-abundance blood proteins were excluded, and 52 secreted and membrane-bound proteins were selected as candidates for the qualification phase.

Our study was designed to simultaneously evaluate candidates for the two clinical needs: (i) differentiation between PCa and negative biopsy, and (ii) discrimination between low-grade (Gleason score (GS) =6) and high-grade (GS ≥8) PCa. We acknowledge that definition of PCa aggressiveness based on GS may not be perfect, however, the correlation between GS and the 20-year survival rate is well established (>70% for GS≤6 and <30% for GS≥8) (29).

### Differential transcriptomics

Genomic alterations in PCa may lead to deregulation of mRNA transcription and protein translation. Since mRNA levels explain only some variation of protein levels (30), mRNA fold changes in tissues and protein fold changes in SP were considered here as independent criteria. To identify candidate genes based on the differential transcriptomics approach, we mined the Cancer Genomics database (www.cbioportal.org) which contained gene expression microarray data in 131 primary PCa tissues and 29 adjacent benign prostate tissues (31), as well as clinical data, such as PSA at diagnosis and GS. The following cut-off criteria were used to select candidates: (i) ≥1.5-fold increased or decreased RNA expression in PCa tissues (N=109) versus adjacent benign prostate tissues (N=29) with *P*-values <0.05 (**Supplemental Table S1**); (ii) ≥1.5-fold increased or decreased expression of mRNA transcripts in PCa tissues with GS =6 (N=41) versus GS ≥8 (N=15) with *P*-values <0.05 (**Supplemental Table S2**); (iii) secreted and membrane-bound proteins based on predicted signal peptides or transmembrane regions (32); and (iv) proteins previously identified in our SP proteome of 3,000 proteins and thus amenable to quantification in SP by SRM assays. Interestingly, a non-coding transcript PCA3 emerged as a top candidate and differentiated between PCa and adjacent benign tissues (6-fold higher expression in PCa with *P* =8×10^−19^, Supplemental Fig. S1). As a result of the differential transcriptomics analysis, we selected 39 candidates.

### Differential proteomics

Shotgun mass spectrometry was used to identify differentially expressed proteins in pools of SP samples (5 samples per pool) from patients with serum PSA ≥4 ng/mL and negative biopsy, low-grade PCa (GS =6) and high-grade PCa (GS ≥8). The rationale for using pooled samples was to reduce the inter-individual variability and increase the likelihood of identifying differences which were consistent between men with a similar diagnosis. A bottom-up proteomic approach and two-dimensional liquid chromatography (2D-LC-MS/MS) followed by the label-free quantification were utilized (30). Proteins were denatured, reduced, alkylated and digested by trypsin. Three process replicates were generated for each pool. Each replicate was fractionated by strong-cation exchange chromatography into 25 fractions, which were analyzed by tandem mass spectrometry (**Supplemental Fig. S2**). Label-free quantification (MaxQuant and Perseus software) was used to prioritize candidates (**Fig. 1C**).

The following cut-off criteria in Perseus were used to select candidates: (i) over- or under-expressed proteins (FDR≤1%, s0=0.22) in high-grade PCa versus negative biopsy (**Supplemental Table S3**); (ii) over- or under-expressed proteins (FDR≤1%, s0=0.23) in low-grade PCa versus negative biopsy (**Supplemental Table S4**); (iii) over- or under-expressed proteins (FDR≤1%, s0=0.27) in high-versus low-grade PCa (**Supplemental Table S5**); (iv) secreted and membrane-bound proteins based on predicted signal peptides or transmembrane regions (32). High-abundance blood serum and testis-, seminal vesicle- and epididymis-specific proteins were excluded. As a result of the differential proteomics analysis, we selected 52 candidates.

### Differential secretomics

Previously, we identified secretomes of a near-normal prostate epithelial cell line RWPE, two androgen-dependent PCa cell lines (LNCaP and VCaP) and five androgen-independent PCa cell lines (PC3, DU145, PPC1, LNCaP-SF and 22Rv1) (33). Here, we hypothesized that the secretome of androgen-independent cell lines may provide candidates elevated at the later stages of PCa or in the more aggressive PCa. We selected 8 most promising candidates which were identified with at least two peptides and were up-regulated ≥2-fold in androgen-independent versus androgen-dependent and normal cell lines based on spectral counting. These 8 candidates (**Supplemental Table S6 and Fig. S3**) were secreted or membrane-bound proteins and were previously identified in the SP proteome.

### Tissue specificity

Aberrant changes in concentration of prostate-specific proteins may indicate the progressing pathological process in the prostate. In fact, the success of PSA is mainly due to its high tissue specificity. Similar to PSA biomarker, leakage of other prostate-specific proteins into blood serum may indicate destruction of prostate-blood barriers due to PCa progression.

To identify proteins with an exclusive or highly restricted expression in prostate tissue, we used the Human Protein Atlas (www.proteinatlas.org) and BioGPS (http://biogps.org) databases. Human Protein Atlas (v. 9) included 12,238 genes with immunohistochemistry-based protein expression profiles in 66 normal human tissues and cells. To identify tissue-specific proteins, we analyzed Human Protein Atlas raw data and ranked proteins according to their tissue-specific expression in 66 human tissues and cells. Proteins with high or medium immunohistochemical staining in prostate, but not in other four tissues of the male urogenital system (seminal vesicles, epididymis, seminiferous tubules and Leydig cells) were finally selected. We also applied a similar strategy to the BioGPS database and identified tissue-specific genes based on the mRNA expression profiles in 84 normal human tissues and cells. In total, we selected 74 proteins with highly specific expression in the prostate, and 48 of these proteins were found in our SP proteome (**Supplemental Table S7**). The list of candidates included 35 secreted and membrane-bound proteins. We also hypothesized that tissue destruction due to PCa progression may result in the elevated amounts of some intracellular proteins in SP and thus retained 13 prostate-specific intracellular proteins.

### Androgen regulation

Physiological role of prostate is highly dependent on androgens and androgen receptor, which also play crucial roles in the development and progression of PCa (34,35). We hypothesized that androgen-regulated proteins might be differentially expressed in SP of low-grade versus high-grade PCa patients. To select androgen-regulated proteins, we reviewed the high-quality datasets of genes with increased expression upon androgen stimulation in LNCaP cell lines or genes with predicted androgen-response elements (**Supplemental Table S8**). We selected 62 androgen-regulated proteins, 20 of which were secreted or membrane-bound proteins present in our SP proteome.

### Development of a multiplex SRM assay for the qualification phase

SRM is a quantitative analytical assay performed with a triple-quadrupole mass spectrometer (36). The assumption is made that the amount of measured proteotypic peptide represents the amount of the protein of interest. With state-of-the-art SRM assays, up to 100 peptides representing medium- and high-abundance proteins (10 ng/mL – 1 mg/mL) can be measured simultaneously in the unfractionated digest of SP, while achieving coefficients of variation under 20%. The number of sample preparation steps prior to SRM measurements should be kept at a minimum, in order to retain high-throughput analysis and minimize variability. Additional fractionation steps, removal of high-abundance proteins or enrichment protocols were thus avoided. In this study, we used our previously published SRM protocols to assay putative protein biomarkers in various biological fluids (37–39). Briefly, Peptide Atlas database (40) was used to select proteotypic peptides for 148 candidates and 11 tissue- and cell-specific control proteins representing seminal vesicles, Cowper’s glands, epididymis, germ cells, Sertoli cells and Leydig cells (**Supplemental Table S9**). SRM assays in SP, however, were developed for only 82 candidate and 11 control proteins (**Supplemental Table S10**). Moderate success rate of SRM assay development could be explained by the low abundance of proteins in SP and the lack of high-quality tryptic peptides. Finally, 93 proteins were assembled into a single multiplexed SRM assay and entered the qualification phase.

### Qualification of candidate biomarkers

In the qualification phase, we measured 77 candidate and 10 control proteins (**Supplemental Fig. S4**) in 13 negative biopsy, 24 low-grade and 14 high-grade age-matched SP samples (6 proteins were excluded after data analysis). SRM areas for each peptide were normalized to a single spiked-in heavy isotopic internal standard of KLK3 protein, and normalized areas were used to calculate concentrations and then diagnostic specificities, sensitivities and ROC areas under the curve (AUC) for each candidate. Proteins were ranked based on their AUCs (**Supplemental Table S11**). Statistical analysis revealed significant up-regulation of 21 proteins (*P* <0.05) in all PCa versus negative biopsy groups. Control proteins, such as MUC6 protein secreted by seminal vesicles, were not significant between groups. Regarding the second clinical need, statistical analysis revealed significant down-regulation of 8 proteins (*P* <0.05) in high-versus low-grade groups. As a result, 29 proteins were selected for the verification phase.

### Upgrade of the SRM assay for the verification phase

To facilitate rigorous verification of top candidates and measure their absolute concentrations in SP, we spiked in heavy isotope-labeled peptides with trypsin-cleavable tags. We also optimized and shortened LC gradient to 30 min, to allow for the measurement of 24 samples in duplicates per day. We then assessed the efficiency of digestion of peptide internal standards by trypsin and revealed a near complete cleavage of quantifying tags. Using shotgun mass spectrometry, we assessed the following post-translational modifications in peptide internal standards after trypsin digestion: cysteine alkylation, methionine oxidation, formation of pyroglutamate of N-terminal glutamine and deamidation of asparagines and glutamines. Using SRM, we quantified the yield of each modification: cysteine alkylation (>99%; 15 peptides), methionine oxidation (~15%; 2 peptides), formation of pyroglutamate of N-terminal glutamine (~50%; 1 peptide) and deamidation of asparagines and glutamines (~20%; 7 peptides). In addition, we evaluated +2 and +3 charge states for 6 peptides and selected both +2 and +3 forms for three peptides. As a result, we included multiple forms of some peptides into the final SRM method (**Supplemental Table S12**). We also fully investigated pre-analytical parameters for 37 peptides using a quantitative multiplex SRM assay. That included LC-SRM injection reproducibility, trypsin digestion reproducibility and the whole process day-to-day reproducibility for three SP samples with different amounts of total protein (35, 67 and 102 mg/mL, **Supplemental Fig. S5**).

### Verification of candidate biomarkers

Top 29 candidates and 8 control proteins moved to the verification phase and were quantified by SRM (**Fig. 2A** and **Supplemental Tables S12-13**). SP samples (N=222) included: 67 negative biopsy samples, 98 low-grade, 38 intermediate-grade and 19 high-grade PCa. SRM areas for each peptide were normalized to the corresponding spiked-in internal standards (**Fig. 2C**), and normalized areas and calibration curves for each protein (**Fig. 2B** and **Supplemental Fig. S6**) were used to calculate protein concentrations in SP (**Fig. 3**). As a result, only TGM4 protein was found significantly up-regulated (3-fold change, Mann–Whitney U test *P* = 0.006, AUC=0.62) in PCa versus negative biopsy samples (**Table 1**). Regarding high-versus low-grade PCa, no proteins were significantly different, while serum PSA revealed significantly higher concentrations (9.4 versus 5.2 ng/mL, *P* =0.0001). Control proteins exclusively expressed and secreted by seminal vesicles (MUC6), epididymis (SG2A1), germ cells (SACA3) and Leydig cells (VTNC) were not differentially expressed. Since analysis of prostate cancer-specific molecules in urine, such as PCA3, may requires normalization to KLK3 mRNA to account for the sample dilution, we also investigated if normalization of protein concentrations by total KLK3 protein in SP would improve the performance of markers (**Supplemental Fig. S7**). Levels of MUC5B, a mucin exclusively expressed and secreted by the Cowper’s glands (41), were significantly lower in PCa (0.4-fold change, *P* =0.006) before, but not after normalization by KLK3 (*P* =0.052). Due to its exclusive expression in the Cowper’s glands and its potentially high variability in SP, MUC5B thus was not considered as a candidate. Even though normalization by KLK3 improved TGM4 AUC from 0.62 (2.9-fold change, *P*=0.003) to 0.64 (3.7-fold change, *P*=0.0003), such increase may not justify the need to measure an additional protein in the clinical lab. Thus, normalization by total SP KLK3 was not further considered in data analysis.

**Fig. 2.**
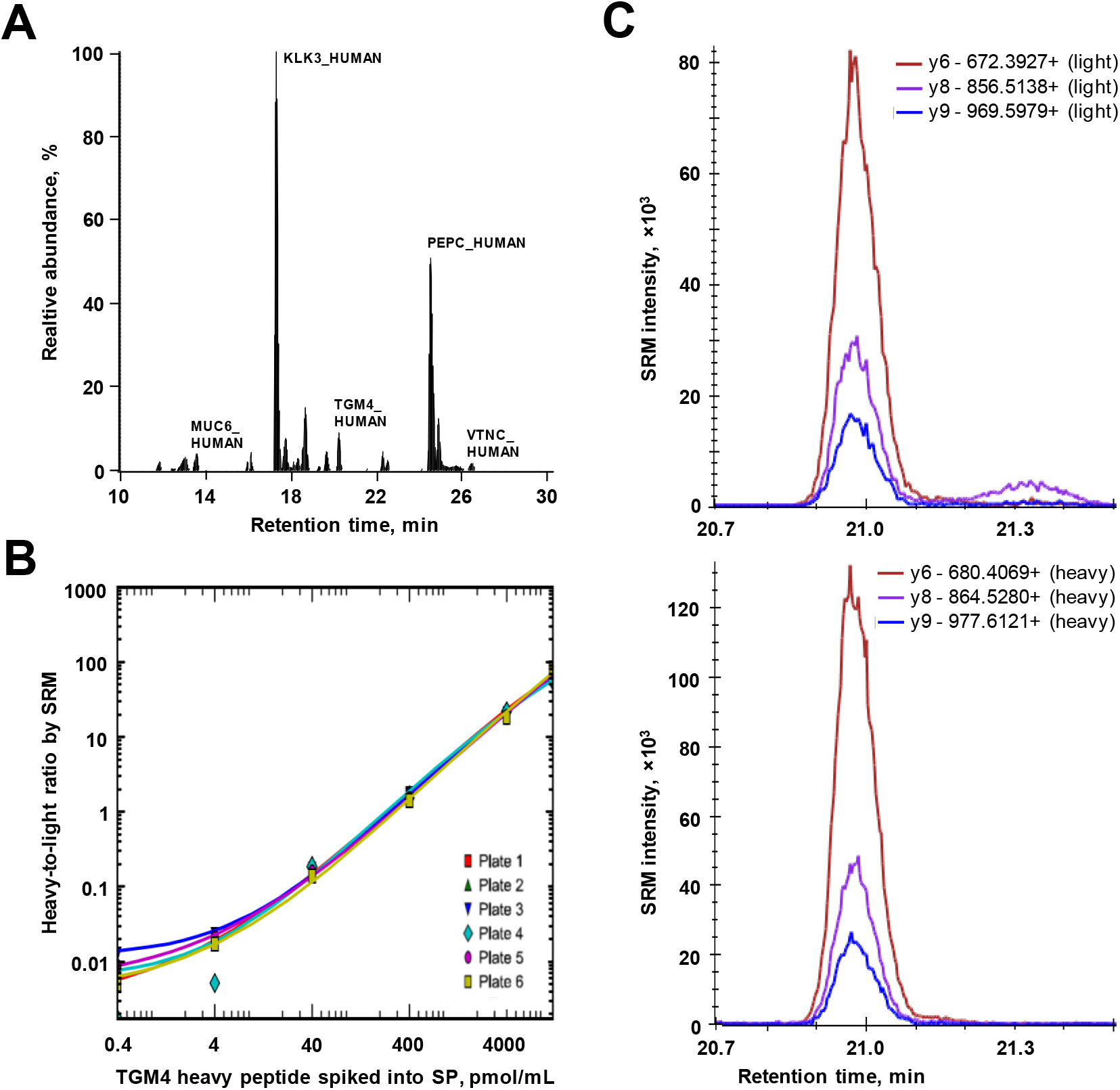
Performance of a multiplex SRM assay in the verification phase. (**A**) Peptides and internal standards representing candidate and control peptides were multiplexed in a single SRM assay within a 30 min LC gradient. (**B**) Representative calibration curves used to quantify TGM4 protein in 67 negative biopsy and 155 PCa SP digests distributed between six 96-well plates. Similar curves were obtained for the rest of proteins (**Supplemental Fig. S6**). Light-to-heavy ratios for TGM4 in each sample were plotted against the corresponding calibration curve, to derive TGM4 concentrations. (**C**) Representative SRM transitions for the light endogenous and heavy IS peptides used to measure light-to-heavy ratio for TGM4 protein in each SP sample.

**Fig. 3.**
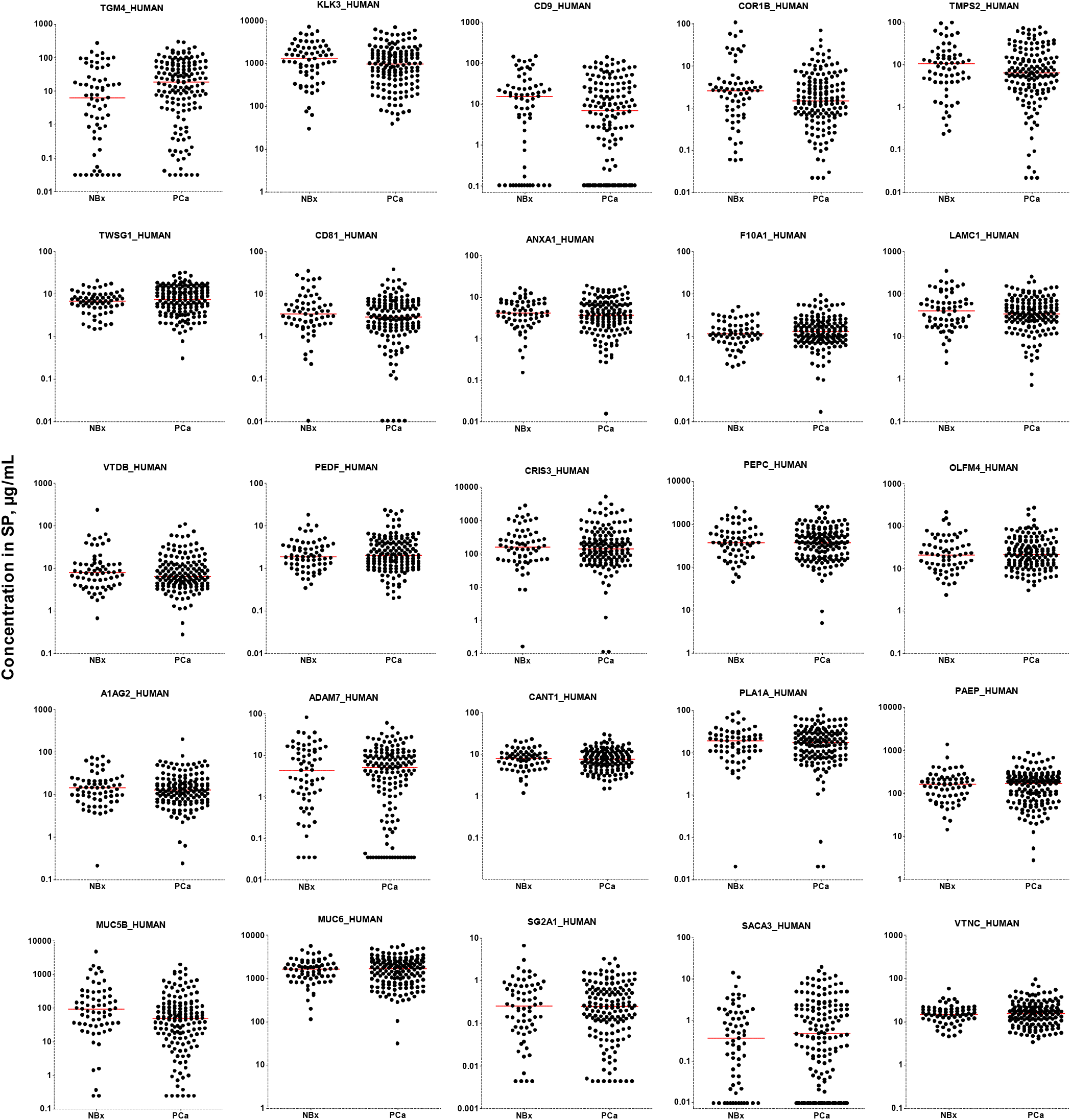
The most promising candidates measured in the verification phase. Using the stable-isotope dilution multiplex SRM assay, 19 candidates and 6 control proteins (KLK3, MUC6, MUC5B, SG2A1, SACA3 and VTNC) were quantified in the negative biopsy (NBx, n=67) and PCa (n=155, all GS) SP samples. Horizontal lines represent median values in SP.

**Table 1.**
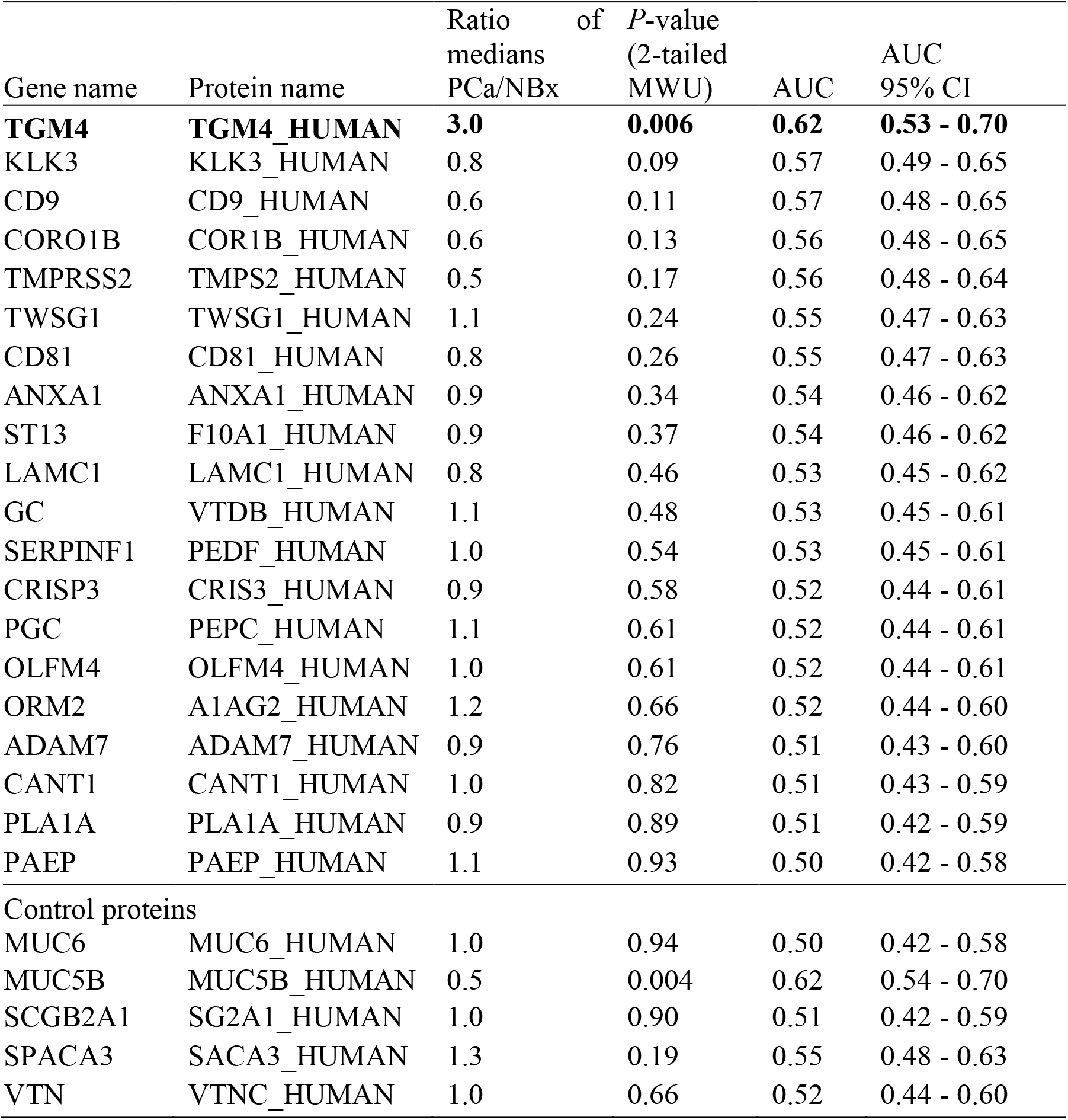
Candidate proteins verified by SRM in 155 PCa and 67 negative biopsy SP samples. Control proteins included proteins exclusively expressed in seminal vesicles, prostate, Cowper’s glands, epididymis, germ cells and Leydig cells. NBx, negative biopsy; MWU, Mann–Whitney U test; AUC, a receiver operating characteristic area under the curve.

| Gene name | Protein name | Ratio of medians PCa/NBx | *P*-value (2-tailed MWU) | AUC | AUC 95% CI |
| --- | --- | --- | --- | --- | --- |
| **TGM4** | **TGM4_HUMAN** | **3.0** | **0.006** | **0.62** | **0.53 - 0.70** |
| KLK3 | KLK3_HUMAN | 0.8 | 0.09 | 0.57 | 0.49 - 0.65 |
| CD9 | CD9_HUMAN | 0.6 | 0.11 | 0.57 | 0.48 - 0.65 |
| CORO1B | COR1B_HUMAN | 0.6 | 0.13 | 0.56 | 0.48 - 0.65 |
| TMPRSS2 | TMPS2_HUMAN | 0.5 | 0.17 | 0.56 | 0.48 - 0.64 |
| TWSG1 | TWSG1_HUMAN | 1.1 | 0.24 | 0.55 | 0.47 - 0.63 |
| CD81 | CD81_HUMAN | 0.8 | 0.26 | 0.55 | 0.47 - 0.63 |
| ANXA1 | ANXA1_HUMAN | 0.9 | 0.34 | 0.54 | 0.46 - 0.62 |
| ST13 | F10A1_HUMAN | 0.9 | 0.37 | 0.54 | 0.46 - 0.62 |
| LAMC1 | LAMC1_HUMAN | 0.8 | 0.46 | 0.53 | 0.45 - 0.62 |
| GC | VTDB_HUMAN | 1.1 | 0.48 | 0.53 | 0.45 - 0.61 |
| SERPINF1 | PEDF_HUMAN | 1.0 | 0.54 | 0.53 | 0.45 - 0.61 |
| CRISP3 | CRIS3_HUMAN | 0.9 | 0.58 | 0.52 | 0.44 - 0.61 |
| PGC | PEPC_HUMAN | 1.1 | 0.61 | 0.52 | 0.44 - 0.61 |
| OLFM4 | OLFM4_HUMAN | 1.0 | 0.61 | 0.52 | 0.44 - 0.61 |
| ORM2 | A1AG2_HUMAN | 1.2 | 0.66 | 0.52 | 0.44 - 0.60 |
| ADAM7 | ADAM7_HUMAN | 0.9 | 0.76 | 0.51 | 0.43 - 0.60 |
| CANT1 | CANT1_HUMAN | 1.0 | 0.82 | 0.51 | 0.43 - 0.59 |
| PLA1A | PLA1A_HUMAN | 0.9 | 0.89 | 0.51 | 0.42 - 0.59 |
| PAEP | PAEP_HUMAN | 1.1 | 0.93 | 0.50 | 0.42 - 0.58 |
| Control proteins |  |  |  |  |  |
| MUC6 | MUC6_HUMAN | 1.0 | 0.94 | 0.50 | 0.42 - 0.58 |
| MUC5B | MUC5B_HUMAN | 0.5 | 0.004 | 0.62 | 0.54 - 0.70 |
| SCGB2A1 | SG2A1_HUMAN | 1.0 | 0.90 | 0.51 | 0.42 - 0.59 |
| SPACA3 | SACA3_HUMAN | 1.3 | 0.19 | 0.55 | 0.48 - 0.63 |
| VTN | VTNC_HUMAN | 1.0 | 0.66 | 0.52 | 0.44 - 0.60 |

### Machine learning analysis to identify multi-variable combinations of markers

To identify combinations of SP proteins that could improve TGM4 performance to differentiate between negative biopsy and prostate cancer patients, or differentiate between high-, intermediate- and low-grade cancers, we employed machine learning algorithms (**Fig. 4A**). A nonlinear Extreme Gradient Boosting (XGBoost) algorithm (19) was selected as the most effective algorithm for obtaining high values for PPV, sensitivity and AUCs after 10×10-fold stratified cross-validation. Relative to other algorithms, XGBoost was better suitable for creating a single strong classifier with a set of weak classifiers and provided better selection of weak features with relative small datasets. Stringent cross-validation was used to reduce over-fitting. Importance of each marker compared to random features was calculated (**Fig. 4B**), and combinations with the highest *F*_05_-scores were selected. AUCs, sensitivities, specificities, PPVs and NPVs were estimated (**Fig. 4C**).

**Fig. 4.**
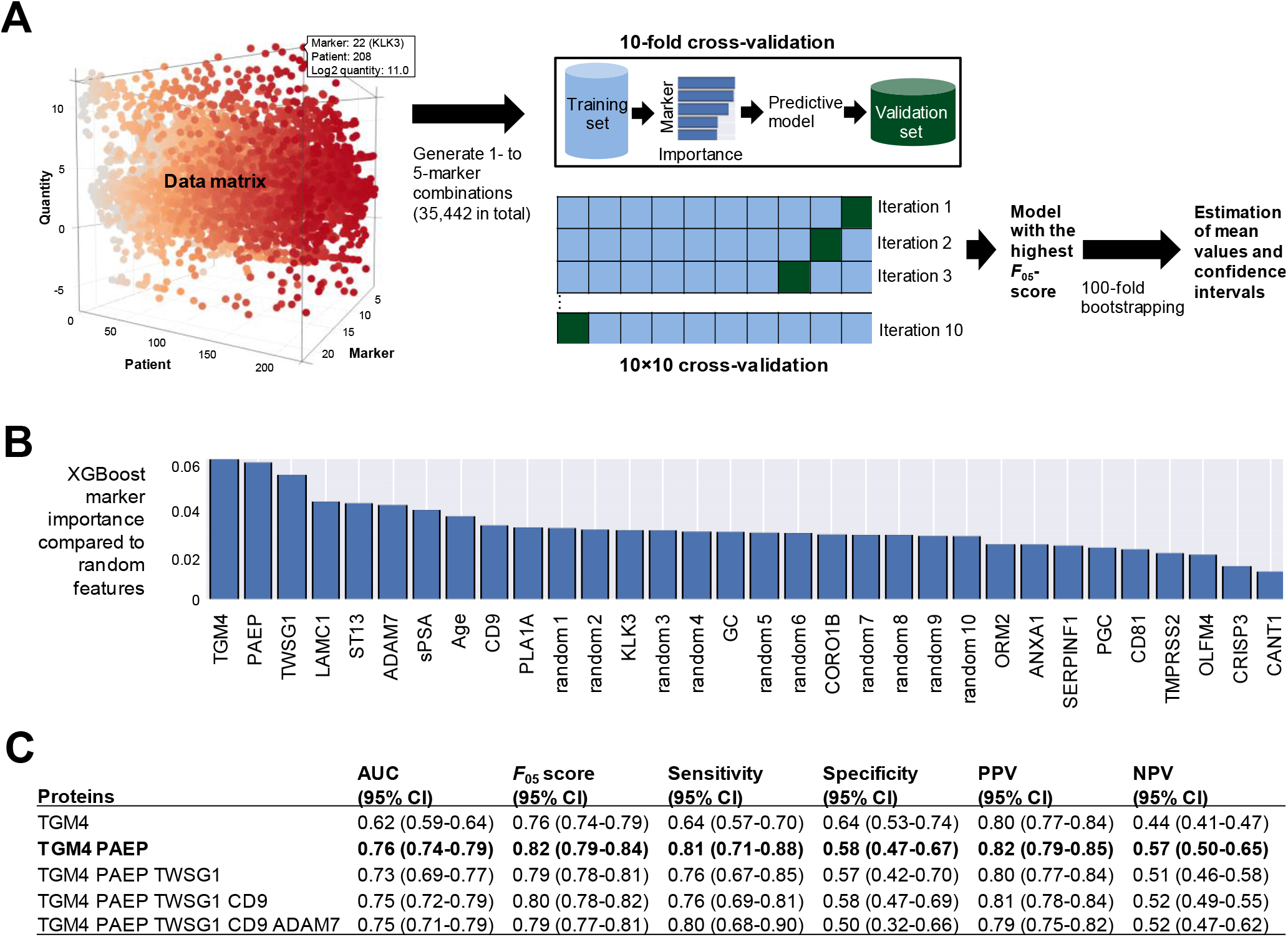
Machine-learning analysis to identify multi-variable combinations of markers. (**A**) Twenty two variables (SP proteins, serum PSA and age) were used to generate all possible 1- to 5- marker combinations (35,442 combinations in total). XGBoost algorithm was used to identify combinations with the highest *F*_05_-measure scores and calculate AUCs, sensitivities, specificities, PPVs and NPVs. Stringent 10×10 cross-validation was applied to reduce over-fitting. Top combinations were verified on the whole dataset of patients to ensure that each potential marker had feature scores higher than a randomly generated feature. Finally, 100-fold bootstrapping was used to estimate mean values for performance metrics and calculate 95% confidence intervals. (**B**) XGBoost marker importance to differentiate between PCa and negative biopsy, as compared to random features. (**C**) Performance of top combinations with 95% confidence intervals estimated using 100-fold bootstrapping. Interestingly, combination of TGM4 with PAEP protein improved AUC and sensitivity to differentiate between negative biopsy and prostate cancer, while additional markers did not further increase AUCs.

Interestingly, combination of TGM4 with PAEP protein improved AUC and sensitivity to differentiate between negative biopsy and prostate cancer, while additional markers did not further increase AUCs. Regarding discrimination of high-grade (Gleason 4+3 and ≥8) versus intermediate-grade (Gleason = 3+4) and low-grade (Gleason 6) cancers, combination of serum PSA with CD9, CORO1B, KLK3 and TMPRSS2 improved AUCs from 0.71 to 0.83, with serum PSA being the strongest classifier (**Supplemental Table S14**). Interestingly, KLK3, PAEP, OLFM4, and PGC presented a combination of SP proteins with the highest AUC = 0.74 (**Supplemental Table S15**). Similar to a 6-peptide signature in the expressed prostatic secretions in combination with serum PSA to discriminate pT2 versus pT3 patients (AUC=0.74) (42), our 4-marker protein signature in SP in combination with serum PSA can be used for a non-invasive detection of high-grade prostate cancer with AUC=0.83.

### Development of TGM4 ELISA

Since our SRM assay (limit of detection 310 ng/mL in SP) was not sensitive enough to measure very low levels of TGM4, we developed in-house ELISA using commercially available sheep and mouse polyclonal anti-TGM4 antibodies and a recombinant human TGM4 (rhTGM4). We determined by immunocapture-SRM assays that endogenous TGM4 from SP was enriched equally well by both antibodies. We also found that unlike mouse antibody, sheep antibody could not efficiently capture rhTGM4. ELISA with a time-resolved fluorescence detection (18) revealed that endogenous TGM4 from SP generated ~3 times higher signal than rhTGM4, and that the mouse-sheep format generated substantially lower background (signal-to-noise 19 for 5 ng/mL) versus sheep-mouse format (signal-to-noise 6). Following that, serial dilutions of endogenous and rhTGM4 and both sheep-mouse and mouse-sheep formats were evaluated. Finally, we selected mouse-sheep format with endogenous TGM4 as a calibrator since such assay format provided the highest signal-to-noise ratio. Limit of blank and limit of detection of our in-house ELISA were 9 and 22 pg/mL, respectively.

### Measurement of TGM4 by ELISA in SP and blood serum

TGM4 was measured by ELISA in 228 SP and 80 blood serum samples (**Fig. 5A, B**). Interestingly, the performance of TGM4 by SRM (3.5-fold change, AUC=0.61, *P*=0.0088) and ELISA (2.9-fold change, AUC=0.62, *P*=0.003) was very similar. In addition, we were able to detect very low levels of TGM4 in blood serum samples (median 120 pg/mL). Unlike SP, serum TGM4 did not differentiate between PCa and negative biopsy. TGM4 levels did not change in prostate inflammation, however, were increased in a much younger population of healthy men (36 versus 65 y.o.). Median TGM4 levels were ~2,000-fold lower in blood than in SP of men with negative biopsy. To the best of our knowledge, our study is the first report on identification and quantification of prostate-specific protein TGM4 in blood serum.

**Fig. 5.**
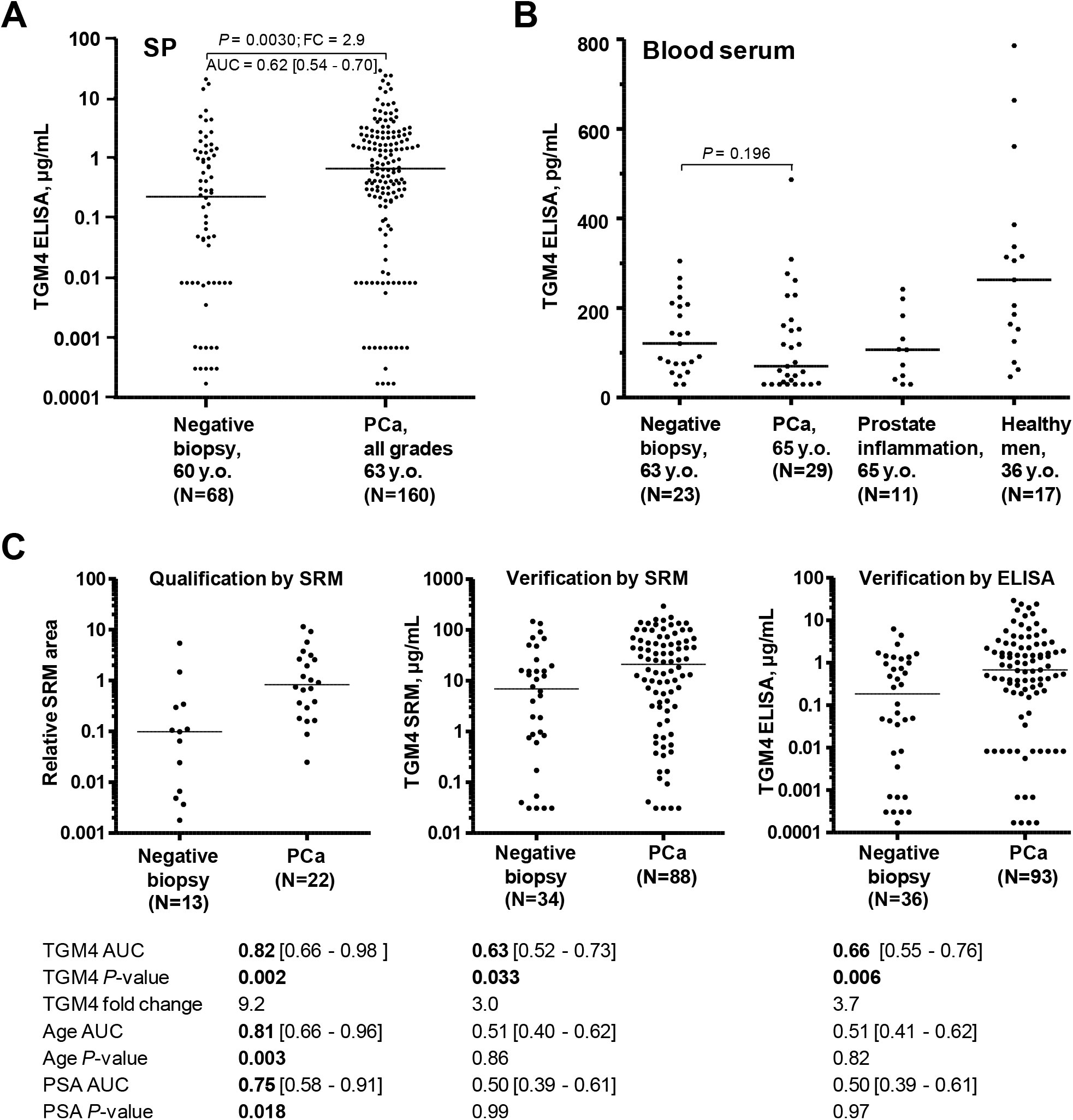
TGM4 performance. Performance of TGM4 as measured by an in-house ELISA in all 228 SP samples (**A**) and 80 blood serum samples (**B**). For comparison, age and serum PSA provided AUCs 0.60 ([95% CI 0.53-0.68]; *P* =0.0105) and 0.56 ([0.48-0.63]; *P* =0.1800) in the same cohorts of patients. (**C**) TGM4 performance in the qualification and independent verification sets for patients with serum PSA ≥4 ng/mL and age ≥50 years old. Qualification included measurements by the label-free SRM, while absolute concentration was measured in the independent verification set by SRM assay with a trypsin-cleavable peptide internal standard. As measured by ELISA, TGM4 had AUC 0.66 to predict PCa on biopsy and outperformed age and serum PSA.

### TGM4 performance in qualification and independent verification sets for patients with age ≥50 y.o. and serum PSA ≥4 ng/mL

Finally, we investigated TGM4 performance in the most relevant group of patients with serum PSA ≥4 ng/mL and age ≥50 y.o. (**Fig. 5C**). Qualification set of samples was measured by a label-free SRM, while an independent verification set (with patients not included into qualification phase) was measured by either SRM with a trypsin-cleavable internal standard or by in-house ELISA. As measured by ELISA, TGM4 protein revealed AUC 0.66 to predict PCa on biopsy and outperformed age (AUC 0.51) and serum PSA (AUC 0.50).

### Correlation of TGM4 concentration with age

We noticed that TGM4 levels in blood serum were higher in samples from younger men versus older patients. To investigate the significance of such trend, we split blood serum samples measured by ELISA into groups of <40 years (N=11), 40-49 (N=11), 50-59 (N=19), 60-69 (N=21) and ≥70 (N=18). As a result, we found a significant difference for TGM4 blood serum levels in these age groups (Kruskal-Wallis *P*=0.0043). The most substantial difference (Dunn’s multiple comparisons test *P* < 0.01) was found for groups with the age <40 y.o. versus >70 y.o. (median levels 206 versus 59 pg/mL). The difference in TGM4 levels for patients in different age groups, as measured by SRM in SP, was not significant (Kruskal-Wallis *P*=0.81). However, TGM4 median levels in SP decreased 3.6-fold between age groups of 40-49 (median 36 μg/mL, N=14) and 70-79 (median 10 μg/mL, N=28). To conclude, the sharp decrease of TGM4 levels after the age of 50 should be carefully considered in future studies and might explain the lack of consensus in the previous studies which identified TGM4 as either up- or down-regulated biomarker of PCa (42–45).

## Discussion

Prostate cancer markers studied up to date include genetic markers, molecular markers such as mRNA, miRNA, proteins and metabolites, and circulating tumor cells. These markers were studied in serum, urine, prostatic secretions, prostate tissues, and cells found in urine, but not in SP. SNP variants of KLK3 and MSMB genes were found to be significantly associated with PCa aggressiveness (46). RNA-based urine biomarkers included PCA3 (8), SPINK1 (10) and the TMPRSS2-ERG fusion (9). Metabolomic profiles in urine and tissues were also investigated in search of prostate cancer biomarkers (47). A few of the more promising protein biomarker candidates included PSA-derived forms and human glandular kallikrein 2 (6,7). None of these other biomarkers has as yet proven to be significantly more sensitive or specific than PSA.

There was little or no reported work on the study of prostate cancer biomarkers in SP. In general, SP is a biological fluid arising from prostate (25% of total SP volume), seminal vesicles (65%), testis and epididymis (10%) and minimal amounts from periurethral glands (48,49). SP acts to support, protect and develop the sperm and is essential for human reproduction. The proteome of SP is as complex as the proteome of serum and contains large amounts of semenogelins, PSA and other high-abundance proteins (14,15). We previously identified more than 3,000 proteins in SP of healthy men and patients with infertility, prostate inflammation and prostate cancer (15,28,50).

Semen and SP are highly relevant biological fluids to search for biomarkers since prostate-secreted proteins are found at much higher concentrations in SP than in serum or urine (13,51). An excellent example of this is PSA. PSA concentration in the SP is 6 orders of magnitude higher than levels found in serum or urine. In fact, much of the work to identify and characterize PSA was originally carried out in SP (12). Since prostatic proteins are more concentrated in semen than in serum or urine, PCa biomarkers might be more easily identified and quantified in SP. We previously discovered male infertility biomarkers in SP and developed a simple 2-biomarker algorithm for the differential diagnosis of male infertility (16,52). Present work might be the largest and the most comprehensive study on PCa biomarkers in SP.

It should be emphasized that semen and SP are unconventional fluids for PCa diagnostics, and that some older patients may have difficulty in providing SP for analysis. However, discussions of our urologists with patients (50 to 75 y.o.) indicated that the vast majority of them were willing and able to provide SP for diagnostic testing, if such test would replace invasive biopsies. Note that older patients (>75 y.o.) are not recommended for screening or biopsy and are usually treated conservatively with active surveillance only (53).

Here, we designed a multi-step biomarker development pipeline which included discovery, qualification and verification phases (54,55). Our study was designed to simultaneously evaluate candidates for two clinical needs: (i) identification of SP biomarkers which could differentiate between PCa and negative biopsy and thus reduce the number of biopsies, and (ii) identification of SP biomarkers which could discriminate between high- and low-grade PCa. Our pipeline was based on quantitative multiplex SRM assays which allowed simultaneous verification of dozens of candidates in hundreds of SP samples within a practical timeline (up to 30-day continuous data acquisition with a mass spectrometer).

We qualified 77 candidates and verified 19 candidates in SP, while TGM4 protein was also verified in blood serum. Many of our candidate proteins have never been previously measured in SP or investigated in the context of PCa, and the molecular function of many proteins is still unknown. Interestingly, levels of numerous prostate-specific proteins previously thoroughly characterized in blood (prostatic acid phosphatase, kallikreins 2 and 3, prostate-specific membrane antigen, beta-microseminoprotein, neuropeptide Y, transmembrane protease serine 2 and others) remained unchanged in SP of PCa patients. In addition, no change was found for the levels of investigated androgen-regulated proteins, except for TGM4.

Multi-variable machine learning analysis provided an interesting combination of TGM4 with a pregnancy-associated endometrial alpha-2 globulin (PAEP). Such 2-marker combination improved detection of PCa on biopsy (AUC=0.76) and should be further investigated in detail.

Previous studies on TGM4 revealed it as a key regulator of invasiveness (56) and cell adhesion (57), and demonstrated association of TGM4 with the epithelial-mesenchymal transition and interaction between cancer and vascular endothelial cells (58). TGM4 was previously suggested as a prostate cancer biomarker, but results were inconsistent and revealed either significant over-expression (43) or under-expression (42,44,45) of TGM4 in PCa versus benign disease, or inconclusive results with the opposite directions based on different assays (59). TGM4 was found down-regulated 1.7-fold in urinary extracellular vesicles of PCa and had AUC 0.58 to diagnose PCa on biopsy (44). Immunohistochemistry with tissue microarrays revealed under-expression of TGM4 in prostate tissues (*P* <0.001) and AUC of 0.81 to detect PCa versus benign disease (44). TGM4 in the urinary extracellular vesicles also differentiated between low- and high-grade PCa with high sensitivity and specificity (*P* <0.001; AUC 0.82). Our present data suggested that TGM4 levels in blood and SP might decrease with age, while TGM4 levels were elevated in PCa versus benign disease. In addition, we observed very high inter-individual variability of TGM4 levels in SP. We thus suggest that combination of these factors (high intra-individual variability, small samples sizes and effect of age and androgen regulation) may explain previous inconsistency regarding the levels of TGM4 in prostate tissues and urine of PCa patients.

In future, TGM4 protein needs to be investigated as a biomarker of distinct subtypes of PCa. For instance, the search of the CamCaP Study Group data (60) revealed that TGM4 mRNA was significantly over-expressed in one of the five clusters (cluster 2, Cambridge cohort of 125 men with primary PCa and matched benign tissues, *P* =0.000011). It should be emphasized that TGM4 has all characteristics of a promising biomarker, such as exclusive prostate tissue specificity, secretion into SP and androgen regulation. Even though demonstrated performance of TGM4 as a single biomarker will unlikely result in its immediate use in the clinic, TGM4 may be further evaluated for the inclusion into emerging multi-biomarker panels. In addition to PCa, TGM4 may be investigated as a biomarker of age, androgen levels and the overall health of prostate.

It should be noted that the major limitation of our study included evaluation of medium- and high-abundance proteins (100 ng/mL - 1 mg/mL) measurable by mass spectrometry in the unfractionated digest of SP. Some promising biomarkers might have been excluded because of their low abundance in SP (<100 ng/mL) and the lack of high-quality SRM assays. In addition, some patients with negative biopsy might have had a missed prostate cancer.

Overall, our study was an ambitious undertaking that involved careful selection of 148 most promising biomarker candidates and quantitative measurements of dozens of proteins in more than two hundred SP samples. Surprisingly, it was only a single protein TGM4 which demonstrated moderate performance to diagnose PCa on biopsy, and none of the candidates differentiated between high- and low-grade PCa. It well may be that “super biomarkers” of PCa may not exist among medium- and high-abundance proteins in SP. Quantification of low-abundance proteins in SP by more sensitive proteomic assays may provide the next generation of PCa biomarkers.

It should also be mentioned that PCa heterogeneity revealed in the recent large-scale genomic studies (61) could hinder identification of true biomarkers. Genomic studies, however, did not find any significant correlation between somatic genomic alterations and PCa aggressiveness (61,62). Thus, true PCa biomarkers may exist only at the downstream epigenetic, proteomic and metabolomic levels. Future proteomic studies should thus take into consideration the distinct genomic subtypes of PCa (61). It well may be that within each unique genomic subtype there will be proteomic signatures which correlate with progression of PCa from low to higher grades. This may facilitate identification of true biomarkers of aggressiveness within the each unique genomic subtype of PCa.

## Acknowledgements

We thank Antoninus Soosaipillai for suggestions on ELISA development, Susan Lau for coordinating collection and storage of clinical samples and Ihor Batruch for assistance with mass spectrometry.

## Author contributions

A.P.D., E.P.D. and K.J. designed the research project. A.P.D. performed all major experiments, analyzed data and wrote the manuscript. P.S. completed shotgun proteomic experiments. M.D. facilitated processing and analysis of SRM data and performed machine learning and statistical analyses. A.D. analyzed transcriptomics data, and T.D.K. assisted with SRM analysis. M.E.H. and K.J. provided clinical samples and clinical expertise.

## Supplemental materials

This article contains supplemental materials.

## Abbreviations

AUC: Area under the curve GS Gleason score
ELISA: Enzyme-linked immunosorbent assay
IQR: Interquartile range
PCa: Prostate cancer
PSA: Prostate-specific antigen
ROC: Receiver operating characteristic
SP: Seminal plasma
SRM: Selected reaction monitoring
TGM4: Protein-glutamine gamma-glutamyltransferase 4

